# No Evidence of Altered Language Laterality in People Who Stutter across Different Brain Imaging Studies of Speech and Language

**DOI:** 10.1101/2024.03.23.586392

**Authors:** Birtan Demirel, Jennifer Chesters, Emily Connally, Patricia Gough, David Ward, Peter Howell, Kate E. Watkins

## Abstract

A long-standing neurobiological explanation of stuttering is the incomplete cerebral dominance theory, which refers to competition between two hemispheres for “dominance” over handedness and speech, causing altered language lateralisation. Renewed interest in these ideas came from brain imaging findings in people who stutter (PWS) of increased activity in the right hemisphere during speech production or of shifts in activity from right to left when fluency increased. Here, we revisited this theory using functional MRI data from children and adults who stutter, and typically fluent speakers (119 participants in total) during four different speech and language tasks: overt sentence reading, overt picture description, covert sentence reading and covert auditory naming. Laterality indices (LIs) were calculated for the frontal and temporal lobes using the LI toolbox running in Statistical Parametric Mapping. We also repeated the analyses with more specific language regions, namely the pars opercularis (Brodmann Area 44) and pars triangularis (Brodmann Area 45). Laterality indices in PWS and typically fluent speakers (TFS) did not differ and Bayesian analyses provided moderate to anecdotal levels of support for the null hypothesis (i.e., no differences in laterality in PWS compared with TFS). The proportions of the PWS and TFS who were left lateralised or had atypical rightwards or bilateral lateralisation did not differ. We found no support for the theory that language laterality is reduced or differs in PWS compared with TFS.

## 1. Introduction

Persistent developmental stuttering is a speech fluency disorder affecting approximately 1% of people worldwide that appears to occur in all languages. The onset of stuttering is typically between ages 2 and 4 years (Yairi & Ambrose, 2013), which corresponds with the rapid development of language skills, such as increase in the average length of utterance (Zackheim & Conture, 2003). The occurrences of stuttering are highly variable both within and across individuals. For example, the frequency of stuttered syllables is affected by different factors, including the complexity of grammar and the length of the planned utterance (Howell et al., 1999; Logan & Conture, 1995; Zackheim & Conture, 2003). Language proficiency is typically unaffected in people who stutter (PWS) (Reilly et al., 2009; Yairi & Ambrose, 1999); PWS know what they want to say, but the flow of speech is disrupted by repetitions, prolongations, and blocks.

Language is perhaps the most lateralised cognitive function: showing primarily left hemisphere involvement for around 96% of right-handers and 70% of left-handers; whilst the rest, 4% of right-handers and 30% of left-handers have what is called atypical lateralisation, indicating either bilateral or primarily right hemisphere involvement (Rasmussen & Milner, 1977). The underlying neurophysiology of stuttering is not yet fully understood, however, one popular neurological explanation of stuttering is the incomplete cerebral dominance theory, which suggests altered patterns of hemispheric specialisation and competition between hemispheres for “dominance” over handedness and speech (Orton, 1927; Travis, 1978). Although scientific interest in the incomplete cerebral dominance theory in stuttering declined over recent decades, it persists as an explanation frequently given by PWS themselves. For example, some PWS describe themselves as being naturally left-handed, but due to the forced use of the right hand, say on school entry, they started to stutter. It is argued that excessive use of the nondominant hand weakens the dominant hemisphere over time. This can cause conflict between hemispheres, leading to altered language lateralisation, which refers to the dominance of one cerebral hemisphere over the other in managing language functions (Orton, 1927).

The first functional neuroimaging meta-analysis in stuttering appeared to support the incomplete cerebral dominance theory by suggesting a tendency towards rightward lateralisation in PWS relative to controls during speech production tasks (Brown et al., 2005). Overactivation in the right inferior frontal cortex is commonly reported in stuttering during speech production tasks, particularly activity of the right frontal operculum and anterior insula (Neef et al., 2018; Watkins et al., 2008) extending to the orbitofrontal cortex (Kell et al., 2009). Moreover, the overactivity in the right hemisphere is described as being “normalised” with speech therapy resulting in typical left lateralised activity (De Nil et al., 2003; Kell et al., 2009; Neumann et al., 2005). Patterns of activity and timing of responses in auditory networks during passive listening tasks also differ between PWS and controls (Beal et al., 2010; 2011; Sato et al., 2011; Weber-Fox & Hampton, 2008). Sato and colleagues (2011) using near-infrared spectroscopy documented diverse laterality in PWS during passive listening to pairs of syllables that included either a phonemic or prosodic contrast, which typically produce left- and right-lateralised patterns of activity respectively in fluent speakers. Notably, none of the adults, school-aged children or pre-school children who stuttered showed significant expected patterns of lateralization for either phonemic or prosodic contrasts, and some showed the reverse pattern.

Even though there are consistent findings of overactivity in the right hemisphere in the functional imaging literature, it still needs to be clarified whether these findings reflect an altered pattern of lateralization. To demonstrate this, it is necessary to statistically compare activity in the two hemispheres during different speech and language tasks. This approaches typically uses a laterality index (LI) calculated using the formula LI = (L-R) / (L+R) to compare the functional activity between two hemispheres. Positive LIs indicate leftwards lateralisation and negative LIs indicate rightwards lateralisation and LIs between -0.2 ≤ LI ≤ 0.2 indicate weak lateralisation, sometimes called “bilateral” (Wilke & Lidzba, 2007). The values entered into the formula for the left and right hemispheres reflect the amount of activity in a given brain region. This is expressed as percent signal change from a baseline or other contrast, or in terms of the number of activated voxels above a certain threshold. This threshold is sometimes referred to as “activated tissue volume,” or it can be the sum of the values of these voxels, which is typically a statistic that reflects the strength of a voxel’s correlation with the task. LIs calculated in this way from brain imaging data can be strongly influenced by individual differences in brain activation and noise levels, the statistical threshold employed, and statistical outliers (Wilke et al., 2006). For example, above a particular threshold, no active voxels might remain in the nondominant hemisphere leading to LI = 1 for the dominant hemisphere, which is biologically implausible. The LI toolbox (Wilke & Lidzba, 2007) running in SPM (Statistical Parametric Mapping: www.fil.ion.ucl.ac.uk/spm/software/spm12/) avoids threshold dependency by calculating LI values across multiple threshold levels and uses a bootstrapping approach to reduce the effect of outliers. The LI toolbox has been successfully employed to measure laterality because of these advantages (Bradshaw et al., 2017).

To address the incomplete cerebral dominance theory in stuttering, we calculated laterality indices for four different speech and language tasks performed using functional MRI in PWS and typically fluent speakers (TFS). Our analyses of these laterality indices were used to target two main questions: (1) Are there any differences between PWS and TFS in functional lateralisation across speech and language tasks? (2) Does functional lateralisation vary across tasks in PWS or TFS or both?

## 2. Methods

### 2.1. Participants

We combined data from a several cohorts of children and adults who stutter who were scanned over the past decade using fMRI during different speech and language tasks. Following exclusion for excessive movement during scanning (details below), useable data were available in 119 participants, 61 PWS (*M*age = 29 years, range = 14-54 years), and 58 TFS (*M*age = 28 years, range = 14-53 years). The groups were not significantly different in terms of age: *t*(117) = 0.60, *p* = 0.55. Chi-squared analysis showed that the groups did not differ in terms of the distribution of handedness (χ2 = 2.23, *p* = 0.13) or gender (χ2 = 2.22, *p* = 0.14). One study with 38 participants (23 PWS, 15 TFS) included only right-handed male participants, therefore we repeated the chi-squared analysis assessing the distribution of handedness after excluding data from this study and again found no difference between groups in terms of the distribution of handedness (χ2 = 3.40, *p* = 0.06). For the PWS group, stuttering ranged in severity from very mild to very severe (9 – 46, median 26) according to the Stuttering Severity Instrument (SSI) (Riley., 1994) At the time of the scan, stuttering severity was assessed as very mild in 8, mild in 11, moderate in 25, severe in 11, and very severe in 6 PWS. The largest subset of participants had performed one of four slightly different versions of overt sentence reading: 56 PWS, and 53 TFS. Smaller subsets had completed tasks involving overt picture description (16 PWS and 18 TFS) and two covert speech tasks involving sentence reading (12 PWS and 12 TFS) and auditory naming (12 PWS and 16 TFS). Table 1 shows the distribution of handedness and sex between groups.

**Table 1.**
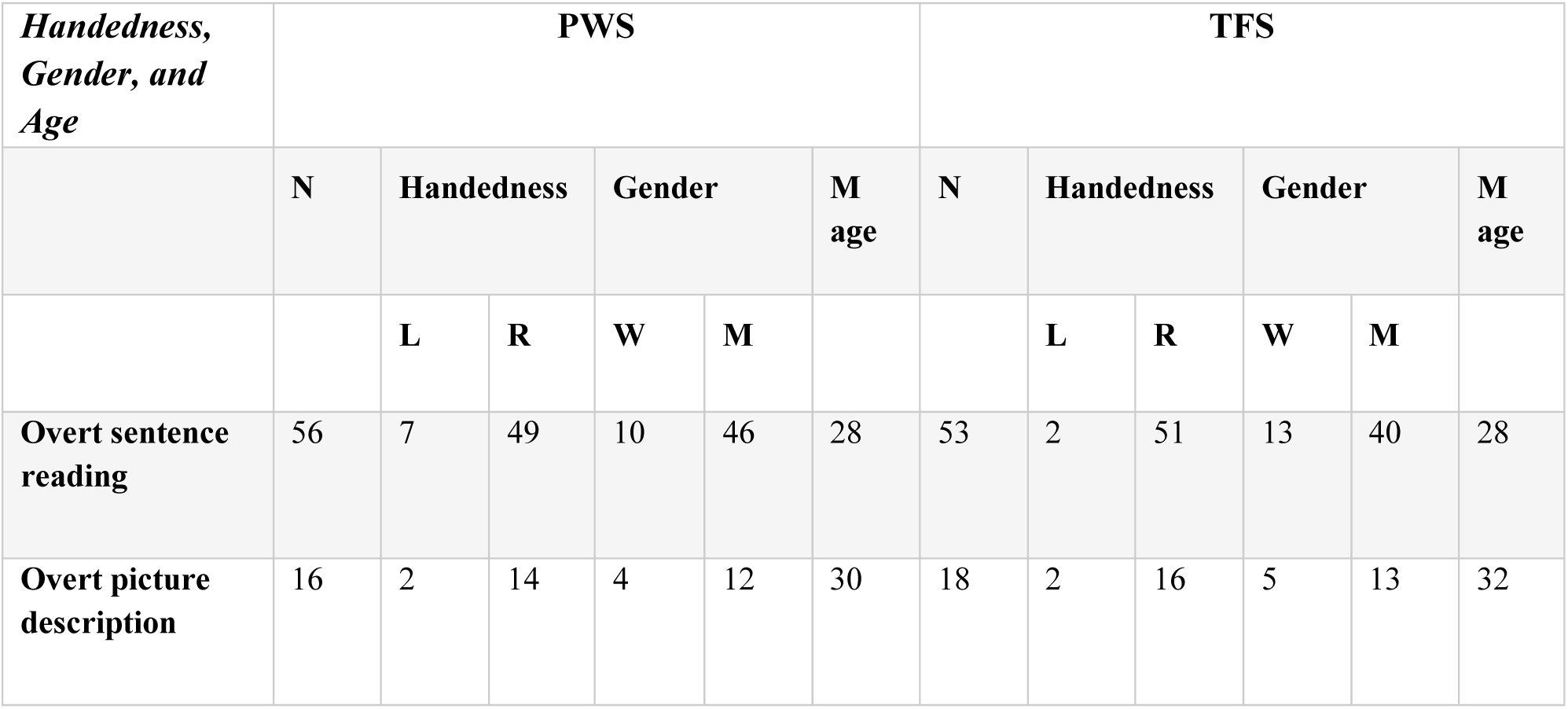
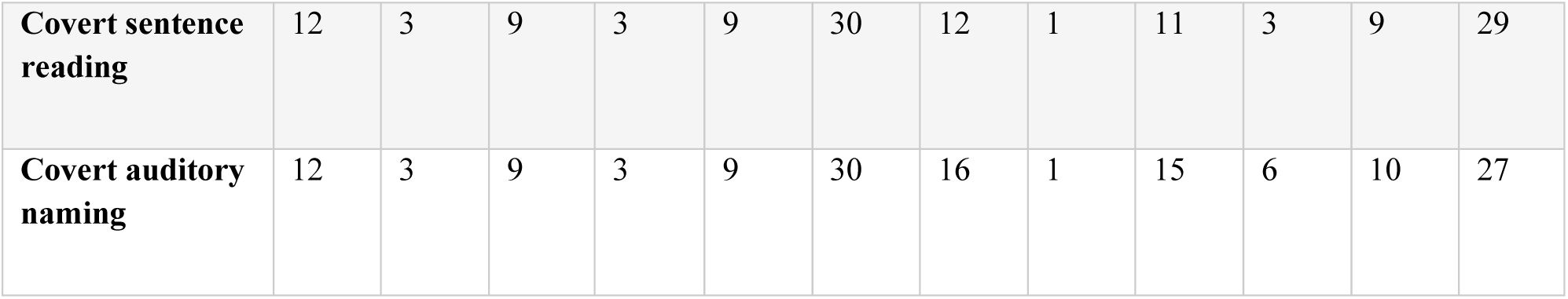
Distribution of handedness and sex of groups across tasks. PWS: people who stutter; TFS: typically fluent speakers; L: Left; R: Right; W: Women; M: Men; Mage: Mean age (in years).

### 2.2. Tasks

The tasks analysed included overt sentence reading, combined from different studies that have been conducted in our laboratories, overt picture description, covert sentence reading, and covert auditory naming tasks (Fig. 1). Overt speech refers to audible production of words/sentences, while covert speech refers to silent production of words/sentences with no articulation. A sparse sampling design (Hall et al., 1999) was used in the overt speaking tasks, in which participants spoke during the silent period between scans. This ensures that the imaging data are not confounded with speech-movement-related artefacts and reduces the possible fluency-enhancing effect of the scanner noise. It also allowed participants to hear the speech they produced. In contrast, continuous acquisition protocols of typical echo-planar imaging were used for the covert speech tasks. Details of the scanners used, and scanning parameters are provided in Supplementary Table 1 with reference to published versions of these studies.

**Figure 1.**
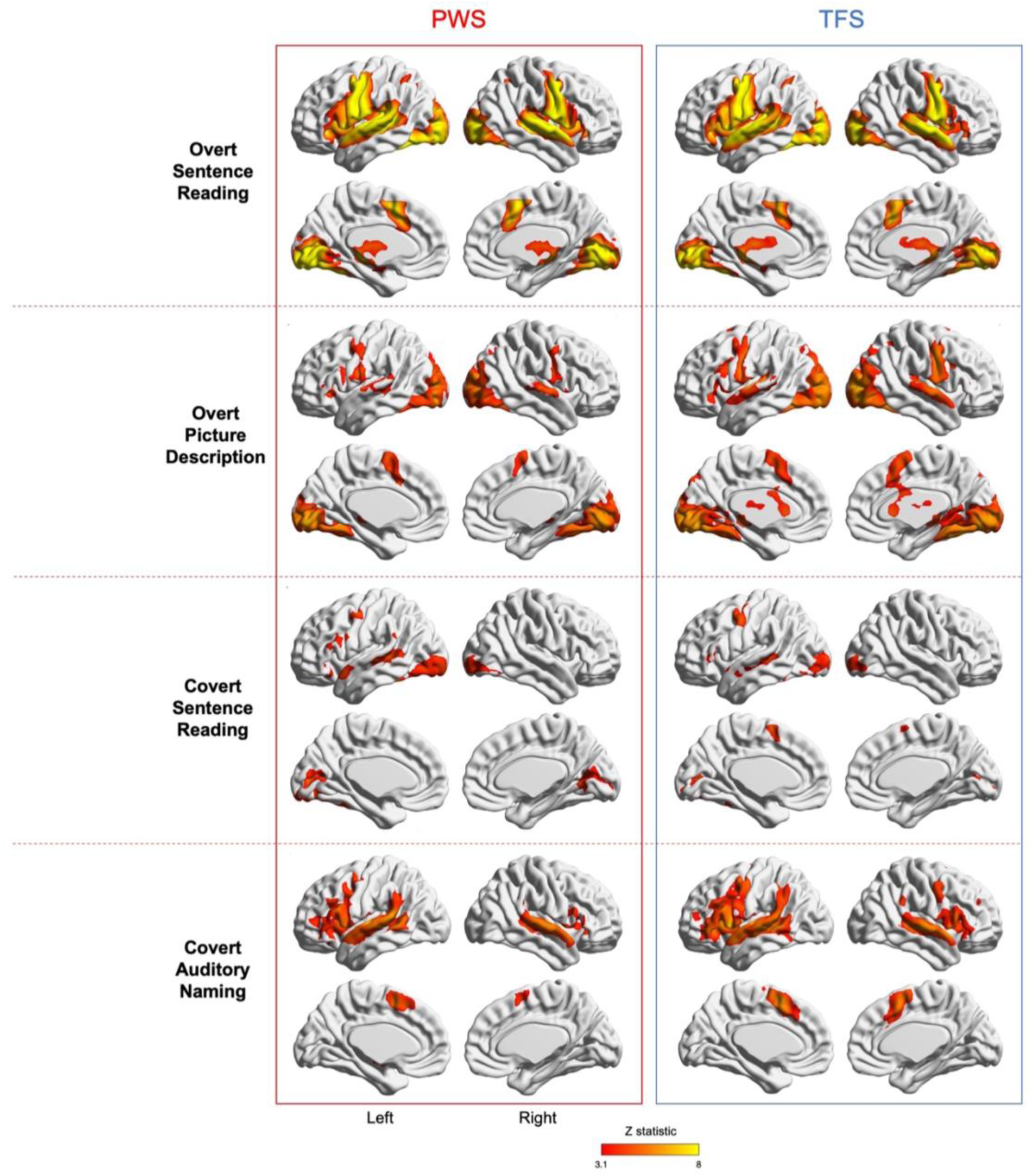
Average brain activity for groups of PWS and TFS during four different tasks. Patterns of brain activity were similar in the two groups. The coloured statistical map is thresholded at Z > 3.1 and p < .05 and overlaid on the MNI-152/Talairach surface template using BrainNet Viewer (Xia et al., 2013). Medial and lateral surfaces of the left and right hemispheres are shown. PWS: People who stutter are on the left of the image; TFS: Typically fluent speakers are on the right.

#### 2.2.1. Overt Sentence Reading

During all versions of the sentence reading task, participants saw a sentence printed in white on a black screen. In some participants (34), a scene corresponding to the sentence was also visible. All participants read the sentence out loud and could hear themselves speaking over headphones during the silent intervals between scan measurements. The baseline conditions consisted of viewing a row of letter X’s that replaced all other letters in a sentence (37 participants), or sentences written in the Farsi alphabet (38 participants), which was unreadable by participants, or a fixation cross (34 participants). Activity during sentence reading was contrasted with the baseline to reveal the brain areas involved in reading aloud, which involves visual word recognition, semantic and syntactic processing, phonological assembly, speech articulation, and monitoring of self-produced speech.

#### 2.2.2. Overt Picture Description

The task was to describe a picture that appeared on the screen using the participant’s own words and spoken out loud. As for overt sentence reading, participants could hear their speech during the silent intervals between scans. The task required conceptualization of the scene, lexical selection and retrieval, phonological and syntactic planning, speech articulation, and monitoring of self-produced speech. A fixation cross was used as a baseline for the picture description task (Connally et al., 2018).

#### 2.2.3. Covert Sentence Reading

Participants read sentences in their imagined speech. In contrast to the overt sentence reading task, reading a sentence covertly does not involve execution of articulatory movements or sensory feedback. This task was contrasted with a row of Xs presented on the screen.

#### 2.2.4. Covert Auditory Naming

Participants heard a short phrase (e.g. “bees make it” over headphones and were asked to generate a word in response (e.g. “honey”) in their imagined speech. The task requires decoding the phonological and semantic information of the auditory input, lexical search and retrieval, and phonological form encoding to generate a word associated with the semantic description (Badcock et al., 2012). A silent baseline and reversed speech baseline conditions were used for the auditory naming task.

### 2.3. fMRI Data Analysis

The fMRI data were analysed at the single-subject level using FEAT (FMRI Expert Analysis Tool part of the FMRIB Software Library; Woolrich et al., 2001). The echo-planar images were motion corrected, smoothed using an 8-mm full-width at half-maximum smoothing kernel, undistorted using fieldmaps where available and registered to the individual’s T1-weighted structural image using boundary-based registration. These individual T1-weighted images were then registered to the MNI-152 template using FMRIB’s non-linear registration tool (FNIRT embedded in FSL). The six motion regressors from the motion correction were included as covariates of no-interest. For continuous imaging sequences (the two covert tasks), the task design was convolved with the double-gamma haemodynamic response function and temporal derivatives were included in the general linear model. For sparse-sampling sequences, convolution with the haemodynamic response function and temporal derivatives were turned off.

Data from five additional scans from the PWS group were excluded from further data analysis due to excessive head movements during scanning. Exclusion criteria were set concerning the mean displacements of head movements being larger than the widest voxel dimension (i.e., > 4 mm) relative to subsequent time points or a warning from the MCFLIRT motion correction tool (Jenkinson et al., 2002) concerning high levels of motion.

### 2.4. Quantifying Laterality Indices

The statistical maps for individual participants were registered to the MNI-152 standard space using their individual T1-weighted structural images (see above) and transferred to the Statistical Parametric Mapping software (http://fil.ion.ucl.ac.uk/spm/) for use with the Laterality toolbox (Wilke et al., 2006). The LI toolbox uses a weighted bootstrapping algorithm to create LI values from the fMRI data by obtaining voxel values for both the left and right hemispheres. First, a vector of voxel values is created from the masked and thresholded input image. Then the algorithm resamples the voxel values 100 times each from the left and right regions of interest (ROIs). All possible LI combinations are calculated from these samples 10,000 times. This procedure is repeated for 20 different threshold intervals (from 0 to the maximum threshold value). Finally, all LIs are plotted on a histogram, and only the central 50% is used as the final LI measure to reduce the influence of outliers. In addition, the LI toolbox produces a weighted mean from the results obtained at different thresholds, which ensures that voxels showing a higher correlation with the task have more impact on the result. Therefore, LI values acquired at a higher threshold are given more weight in the final LI-weighted mean value.

ROIs were chosen based on masks of the frontal and temporal lobes, as these include the regions typically activated by the language tasks. Activity from voxels in the medial wall was excluded from the LI calculations since it is difficult to determine the hemispheric origin of activity at this location due to smoothing (standard exclusion mask: midline -5 mm). In separate analyses, we also measured the LI values in smaller ROIs, namely the pars opercularis (BA44) and pars triangularis (BA45) of the inferior frontal gyrus, to provide a more granular analysis. Masks were generated using the Juelich atlas within FSL software, for the BA44 and BA45 brain regions with a threshold set at 30%.

### 2.5. Statistical Analysis

Initial inspection of the data suggested no group differences in terms of laterality indices (see Fig. 2 below); therefore, we selected a Bayesian approach in order to weigh the evidence in favour of the null relative to the alternative hypothesis. We performed Bayesian t-tests to address our first research question to test whether there are differences in LI-weighted values between PWS and the TFS group for four different speech and language tasks. We reported the Bayesian Factor 01 value, which refers to the ratio of the ‘likelihood of data given the null hypothesis / likelihood of data given the alternative hypothesis’. In alignment with Jeffreys’ guidelines (Jeffreys, 1939), we considered a Bayes factor falling within the range of 1 to 3 as indicative of anecdotal evidential support, while a Bayes factor ranging from 3 to 10 signifies a moderate level of evidential support. A Bayes factor surpassing 10, on the other hand, indicates robust evidential support. We also categorized datasets as showing typical patterns of leftwards laterality (> 0.2) compared with atypical (< -0.2 right or bilateral, -0.2 to 0.2 LIs), to determine whether the distribution of laterality in individuals differed between groups using chi-squared analysis.

**Figure 2:**
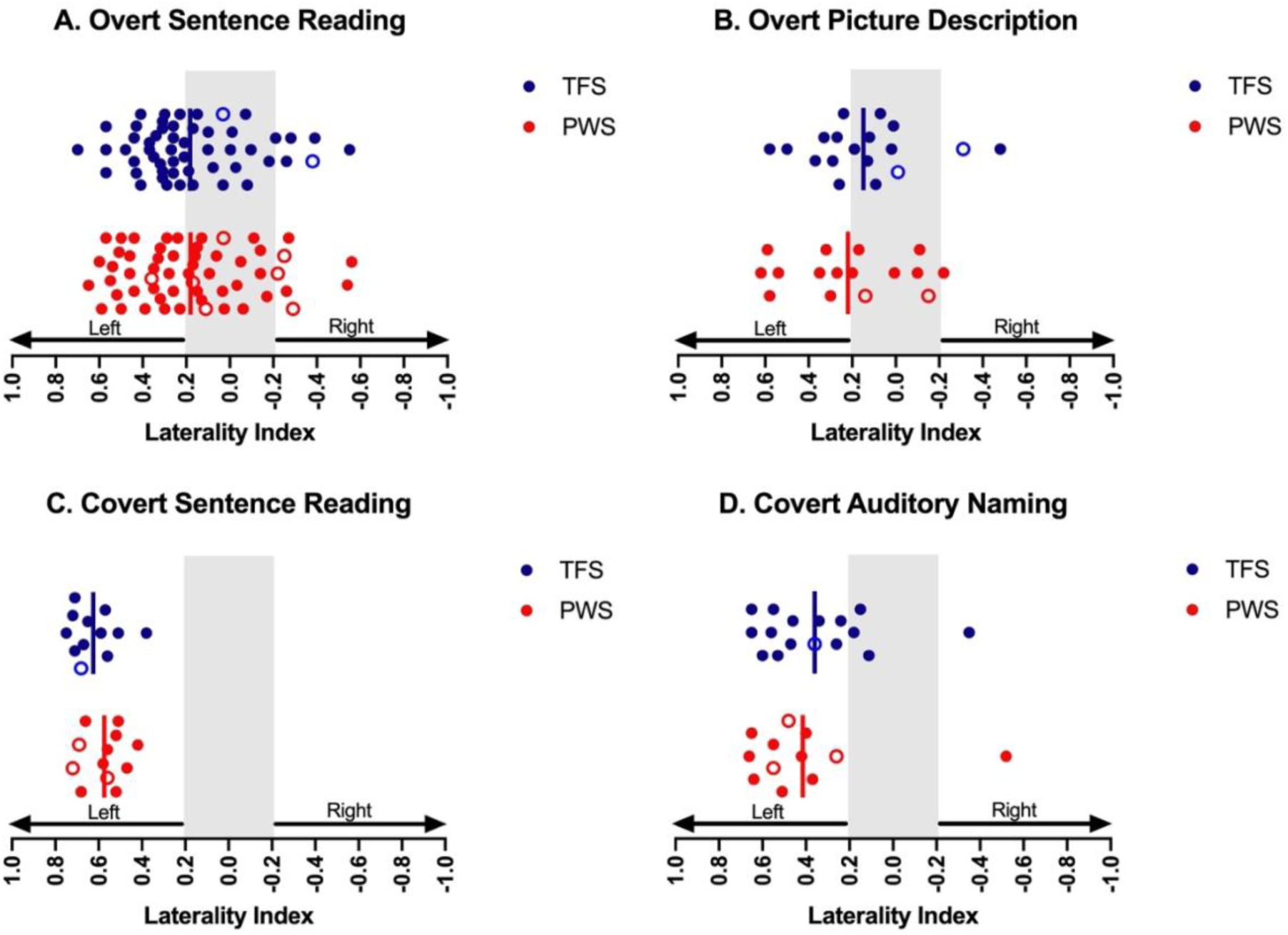
Laterality indices in PWS and TFS based on activity in the frontal lobes. Solid vertical lines represent group means. The grey area represents LI values between -0.2 and 0.2, which are considered not lateralised. PWS: People who stutter are the red circles; TFS: Typically fluent speakers are the blue circles. The open circles indicate left-handedness in both groups.

To compare laterality indices among the different tasks, linear mixed model regression analyses were performed. All Bayesian and chi-squared tests were carried out using the software JASP (JASP Team, 2022) and we used RStudio (RStudio Team, 2022) to perform linear mixed model regression analyses.

## 3. Results

### 3.1. Brain activity during tasks: Group Averages

We looked at the brain activity of each group relative to the baseline condition for each task based on a cluster-forming threshold of Z > 3.1 and an extent threshold of p < .05 (corrected for family-wise error). Both groups showed a broad range of activity in brain regions expected to be activated by speech and language (Fig. 1; Table 2).

**Table 2.** Laterality indices across tasks for the frontal and temporal lobes, BA44 and BA45. LI: Laterality indices’ weighted mean; SEM: Standard Error of Mean; Typ: Participants who show typical pattern of leftwards laterality; Atyp: Participants who show right or bilateral laterality.

| *Region x Task* | PWS |  |  |  | TFS |  |  |  |
| --- | --- | --- | --- | --- | --- | --- | --- | --- |
|  | N | Mean LI + SEM | Typ | Atyp | N | Mean LI + SEM | Typ | Atyp |
| *Frontal lobes* |  |  |  |  |  |  |  |  |
| Overt sentence reading | 56 | 0.18 ± 0.03 | 28 | 28 | 53 | 0.18 ± 0.03 | 31 | 22 |
| Overt picture description | 16 | 0.21 ± 0.07 | 9 | 7 | 18 | 0.14 ± 0.06 | 8 | 10 |
| Covert sentence reading | 12 | 0.57 ± 0.02 | 12 | 0 | 12 | 0.62 ± 0.03 | 12 | 0 |
| Covert auditory naming | 12 | 0.41 ± 0.09 | 10 | 2 | 16 | 0.36 ± 0.06 | 12 | 4 |
| *Temporal lobes* |  |  |  |  |  |  |  |  |
| Overt sentence reading | 56 | 0.12 ± 0.04 | 29 | 27 | 53 | 0.13 ± 0.04 | 24 | 29 |
| Overt picture description | 16 | -0.06 ± 0.06 | 4 | 12 | 18 | 0.05 ± 0.05 | 6 | 12 |
| Covert sentence reading | 12 | 0.56 ± 0.06 | 12 | 0 | 12 | 0.58 ± 0.04 | 12 | 0 |
| Covert auditory naming | 12 | 0.23 ± 0.08 | 9 | 3 | 16 | 0.24 ± 0.05 | 11 | 5 |
| *BA 44* |  |  |  |  |  |  |  |  |
| Overt sentence reading | 56 | 0.18 ± 0.05 | 30 | 26 | 53 | 0.20 ± 0.05 | 28 | 25 |
| Overt picture description | 16 | 0.40 ± 0.07 | 12 | 4 | 18 | 0.34 ± 0.07 | 13 | 5 |
| Covert sentence reading | 12 | 0.59 ± 0.03 | 12 | 0 | 12 | 0.51 ± 0.09 | 11 | 1 |
| Covert auditory naming | 12 | 0.50 ± 0.08 | 11 | 1 | 16 | 0.41 ± 0.06 | 14 | 2 |
| *BA45* |  |  |  |  |  |  |  |  |
| Overt sentence reading | 56 | 0.04 ± 0.06 | 24 | 32 | 53 | 0.13 ± 0.05 | 25 | 28 |
| Overt picture description | 16 | 0.34 ± 0.11 | 12 | 4 | 18 | 0.19 ± 0.09 | 10 | 8 |
| Covert sentence reading | 12 | 0.51 ± 0.09 | 11 | 1 | 12 | 0.45 ± 0.11 | 10 | 2 |
| Covert auditory naming | 12 | 0.43 ± 0.09 | 10 | 2 | 16 | 0.40 ± 0.10 | 15 | 1 |

During the overt sentence reading task, large portions of the primary motor cortex, premotor cortex, presupplementary motor area extending to anterior cingulate cortex, superior temporal lobe, the thalamus, ventral occipito-temporal cortex, and lobule VI of the cerebellum were significantly activated bilaterally in the PWS group but somewhat more extensively on the left. The TFS group exhibited strikingly similar brain activity patterns to the PWS group.

The overt picture description task evoked significant activity in premotor and primary motor cortex, the presupplementary motor area, superior temporal gyrus, thalamus, ventral occipito-temporal cortex, and the anterior lobe of the cerebellum, all bilaterally with additional activation of the left inferior frontal gyrus in both groups that was more extensive in the TFS group.

The covert sentence reading task evoked significant activity in inferior frontal cortex, primary motor cortex (at the level of the face representation), superior temporal gyrus, and mid fusiform gyrus in the left hemisphere, and in lateral and medial occipital cortex bilaterally in both groups. There was additional activity in the cerebellar vermis in PWS and presupplementary motor area in the TFS group.

During the covert auditory naming task, there was significant activity in the inferior frontal gyrus bilaterally but more extensively on the left, the presupplementary motor area, the superior temporal gyrus and dorsal striatum bilaterally, and in the right anterior and posterior lobes of the cerebellum in both groups. In the PWS, there was significant activity in the primary motor cortex on the left and this was seen bilaterally in the TFS group; TFS also had activity in the left anterior lobe of the cerebellum.

### 3.2. LIs based on task-evoked activity in the frontal lobes

#### 3.2.1. Overt Sentence Reading Task

The results of our LI calculations for the frontal lobe masks for the sentence reading task are shown in Table 2 and Fig. 2. The sentence reading task was completed by the largest number of participants combined across four different versions of the task. A Bayesian independent samples t-test (two-sided) revealed a Bayes factor of 4.92, indicating moderate evidence in support of the null hypothesis (LIs are similar between PWS and TFS groups) rather than the alternative hypothesis (LIs differ between PWS and TFS). Chi-squared statistics found that the number of individuals in each group who showed the typical pattern of leftwards laterality compared with atypical (right or bilateral LIs) did not differ (χ^2^ = 0.79, *p* = 0.37; see Table 2). It is worth noting that the mean LI for each group falls in the range considered indicative of weak laterality or bilateral (Table 2), indicating that the tasks is not reliably strongly lateralized, though individual values ranged from -0.56 to 0.65 in PWS and -0.55 to 0.70 in TFS.

#### 3.2.2. Overt Picture Description, Covert Sentence Reading, Covert Auditory Naming

In different subsets of the overall sample, we also analysed LIs for the overt picture description task, covert sentence reading task, and covert auditory naming task (Table 2, Fig. 2). Bayesian independent samples t-test analyses revealed Bayes factors of 2.41, 1.55, and 2.56, respectively for these tasks, indicating anecdotal evidence in support of the null hypothesis (no group differences for each task). On the basis of these findings in smaller subsets of the sample, we can neither reject nor support the null hypothesis. Chi-squared analyses found no group differences regarding the number of typically or atypically lateralised individuals in the overt picture description task (χ^2^ = 0.47, *p* = 0.49), covert sentence reading task (all participants were left lateralised) and covert auditory naming task (χ^2^ = 0.28, *p* = 0.59).

#### 3.2.3. Laterality Effect of Different Tasks

Inspection of Fig. 2 indicates that, for the frontal lobes, the two covert tasks (covert sentence reading and auditory naming) were more robustly lateralized at both the group and the individual level relative to the pattern of lateralization seen for the overt tasks (overt sentence reading and picture description). To test this quantitatively, we employed linear mixed models with overt tasks as a fixed effect. Group (PWS and TFS) was entered into the model as interactions for this fixed effect with intercepts of subjects as our random effect. The model revealed that LIs were significantly more left lateralised for covert tasks (N = 51, LI = 0.49, *SE* = 0.03) compared with overt tasks (N = 143, LI = 0.18, *SE* = 0.02) (β = -0.39, *t* = -6.33, *p* < 0.001).

There were no significant interactions with the group.

### 3.3. LIs based on task-evoked activity in the Temporal Lobes

#### 3.3.1. Overt Sentence Reading

For the temporal lobe masks, a Bayesian independent samples t-test revealed a Bayes factor of 4.87, indicating moderate evidence in support of the null hypothesis. The chi-squared analysis gave the same result as the analysis of frontal lobes and verified that the groups did not differ regarding the number of typically or atypically lateralised individuals (χ^2^ = 0.46, *p* = 0.50). As above, the mean LIs indicate this is not a robustly lateralized task at the group level, though again the range of values indicates considerable variation among individuals (Fig. 3; Table 2).

**Figure 3.**
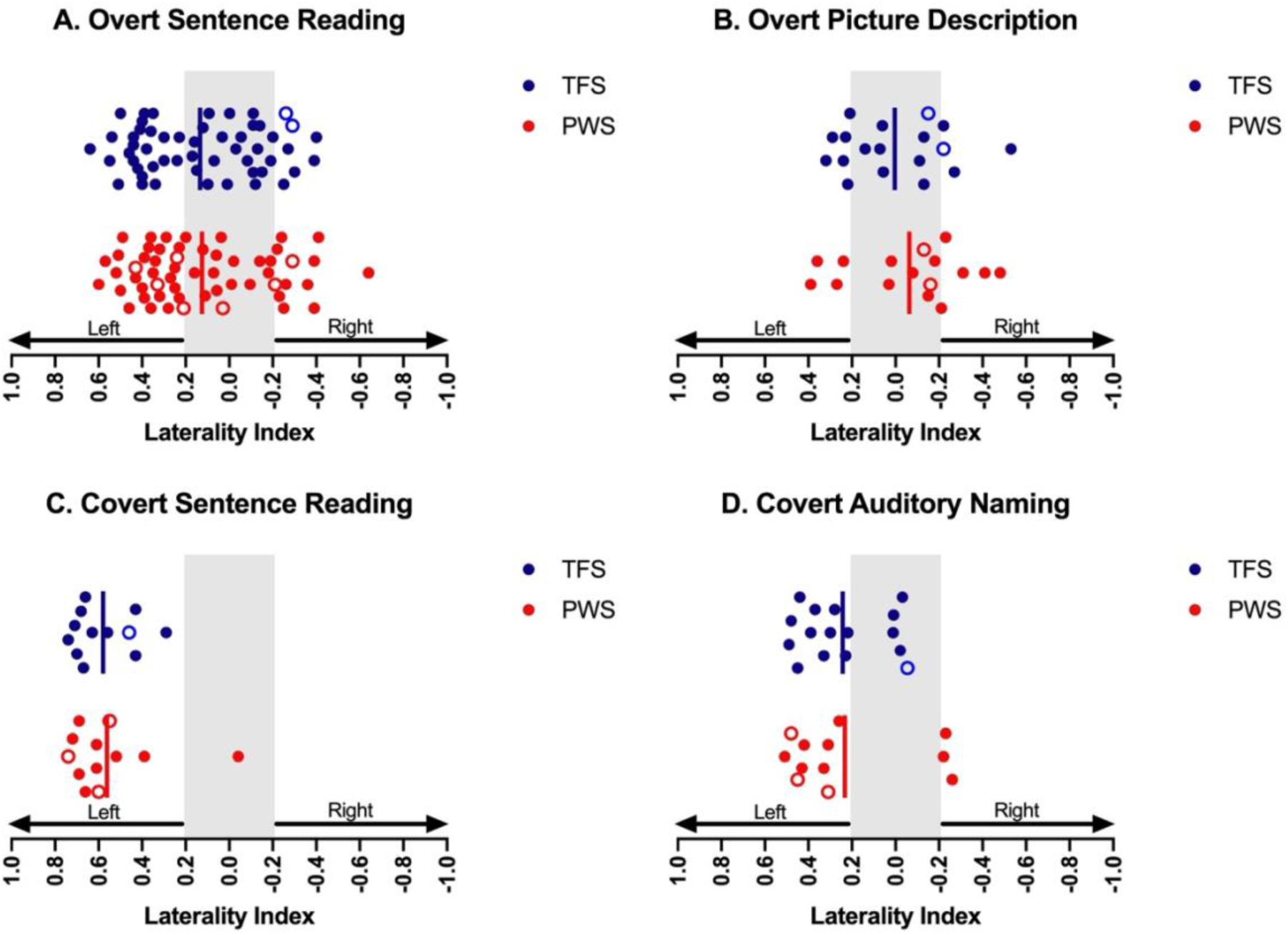
Laterality indices in PWS and TFS based on activity in the temporal lobes (see Fig. 2 for details).

#### 3.3.2. Overt Picture Description, Covert Sentence Reading, Covert Auditory Naming

For the temporal lobe masks, we examined LIs for the overt picture description task, covert sentence reading task, and covert auditory naming task (Fig. 3; Table 2). Bayesian independent samples t-tests found no support for the alternative hypothesis with Bayes factors of 2.36, 2.61, and 2.80, respectively, indicating anecdotal evidence in support of the null hypothesis. Furthermore, chi-squared analyses found the same results as for the frontal lobe analyses, which confirmed that the groups did not differ in terms of the number of typically or atypically lateralised individuals in the overt picture description task (χ^2^ = 0.28, *p* = 0.9), covert sentence reading task (χ^2^ = 1.04, *p* = 0.30) and covert auditory naming task (χ^2^ = 0.13, *p* = 0.72) (see Table 2).

#### 3.3.3. Laterality Effect of Different Tasks

The same linear mixed model analysis as above was performed on the data from the temporal lobe masks to test differences in lateralisation between covert and overt tasks (Fig. 3). Similarly to the above, the model revealed that the covert tasks (N = 51, LI = 0.40, *SE* = 0.04) showed significantly more robust left lateralisation than overt tasks (N = 143, LI = 0.09, *SE* = 0.02) (β = -0.30, *t* = -4.32, *p* < 0.001).

### 3.4 LIs for task-evoked activity in Pars Opercularis (BA44) and Pars Triangularis (BA45)

We analysed the LI values within more focal regions of interest, specifically the pars opercularis (BA44) and pars triangularis (BA45) of the inferior frontal gyrus (Table 2). This approach was adopted to facilitate a more detailed examination since these regions in the left hemisphere correspond to “Broca’s Area” and are usually robustly activated by speech and language tasks. Again, we found similar results suggesting no differences between groups in LIs (see Supplementary figures). For BA44, data from the overt sentence reading task (which has the largest sample size) provided moderate evidence in favour of the null hypothesis with the Bayes Factor (BF) of 4.82. For the other tasks, we found only anecdotal evidence in favour of the null: overt picture description (BF = 2.64), covert sentence reading (BF = 2.24), and covert auditory naming (BF = 2.05). For BA45, we only found anecdotal evidence in favour of the null hypothesis for each of the four tasks; overt sentence reading (BF = 2.91), overt picture description (BF = 2.08), covert sentence reading (BF = 2.49) and covert auditory naming tasks (BF = 2.78).

Chi-squared analyses also revealed no significant differences between the groups in terms of the number of participants in each group who were typically or atypically lateralized: BA44 in overt sentence reading (χ^2^ = 0.006, *p* = 0.93), overt picture description (χ^2^ = 0.03, *p* = 0.85), covert sentence reading (χ^2^ = 1.04, *p* = 0.30), covert auditory naming (χ^2^ = 2.70, *p* = 0.10); BA45 in overt sentence reading (χ^2^ = 0.20, *p* = 0.65), overt picture description (χ^2^ = 1.40, *p* = 0.23), covert sentence reading (χ^2^ = 0.38, *p* = 0.53), and covert auditory naming task (χ^2^ = 0.10, *p* = 0.75) tasks (see Table 2).

## 4. Discussion

Since the pioneering findings of Paul Broca, it is well documented that most people rely more on their left hemisphere than their right to use language. In the current study, we investigated the theory as to whether PWS have reduced hemispheric specialisation compared with typically fluent speakers. We looked at data obtained across different language and speech tasks, namely overt sentence reading, overt picture description, covert sentence reading, and covert auditory naming that we obtained in different functional MRI studies. Pooling data across these different versions allowed us to investigate one task (overt sentence reading) in a relatively large sample of 56 PWS compared with 53 TFS. We analysed LI-weighted means for task-evoked activity in frontal and temporal lobes and repeated the same analyses for portions of the inferior frontal gyrus involved in language processing, namely the pars opercularis (BA44) and pars triangularis (BA45) regions. We did not exclude stuttering epochs from the overt speaking data in this analysis since it is possible that stuttering evokes right hemisphere activity. The inclusion of stuttering epochs therefore made it more likely we would detect a rightwards pattern of lateralisation in people who stutter should one exist.

Our main findings are as follows: (1) there was no evidence in support of the idea that PWS are differently lateralized relative to people who are typically fluent and some anecdotal and moderate evidence in support of the idea that they are equally lateralized; (2) leftwards lateralization was most robustly observed for the covert tasks relative to the overt tasks and this effect was seen in both groups.

### 4.1. No Differences in Laterality between PWS and TFS

The idea that people who stutter are differently lateralized for language has a long history dating back to Samuel Orton (1927), who first proposed this idea, and it is a persistent “urban” myth believed by many individuals with lived experience of stuttering. The support for this myth comes from two main sources of information: (i) it has been suggested that left-handedness may be a risk factor for stuttering and that it occurs more frequently in PWS than in the general population (Howell et al., 2007; see a recent meta-analysis: Papadatou-Pastou et al; 2023), which is often assumed to be associated with more rightward lateralization for language in the brain (Orton, 1927; Travis, 1978); (ii) some early observations and neuroimaging studies proposed atypical hemispheric activity during stuttered-speech that shifted to the left when individuals spoke fluently (Fox et al., 1996; Jones, 1966; Neumann et al., 2005; Kell et al., 2009).

First, our groups of people who stutter and typically fluent speakers did not differ in the distribution of left and right-handedness, while it is important to note that one of the studies with 38 participants (23 PWS, 15 TFS) recruited only right-handed male participants. Nevertheless, the distribution of handedness between our groups remained stable, even when data from that study were excluded from our handedness analysis. Therefore, our data is consistent with evidence indicating that stuttering is independent from handedness (Records et al., 1977; Salihović & Sinanović, 2000; Webster & Poulos, 1987). We find no support for the idea that PWS are more likely to be left-handed than typically fluent speakers, nor that their handedness alters the typical pattern of left-hemispheric cerebral dominance for speech and language.

In our main findings, we found that PWS and TFS show equivalent levels of language lateralization across a range of tasks. The means for groups of PWS and TFS were very similar, and they had very similar distributions of participants categorised as typically (left) or atypically (right or bilaterally) lateralised. This pattern of results was seen for our large region of interest analyses, where we compared task-evoked activity in the frontal and temporal lobes, as well as in our analyses of more focused regions of interest in the inferior frontal cortex. The results of our statistical analyses found no support for the idea that PWS are differently lateralized on average relative to people who are fluent speakers, in fact, in some cases we found moderate support for the hypothesis that they are not differently lateralized. These results are compatible with a magnetoencephalography (MEG) study which compared language lateralisation in preschool children who stutter and controls during picture naming tasks (Sowman et al., 2014). The authors reported that the language was mostly left lateralised in both groups over frontal, temporal and parietal regions without significant differences between groups.

On the other hand, several studies reported additional right hemisphere activity in PWS compared with controls during speech production (Kell et al., 2009; Neef et al., 2018; Neumann et al., 2005) and this right hemisphere overactivity in the inferior frontal cortex or frontal operculum was described as one of the neurophysiological “signatures” of stuttering (Brown et al., 2005). For example, a recent functional imaging study revealed greater activity over the right hemisphere of PWS compared with TFS during both covert speech and covert humming conditions (Neef et al., 2018). This overactivity was interpreted as an overly active global response suppression mechanism mediated via the hyper-direct pathway to the subthalamic nucleus rather than reflecting reorganisation of function to the right hemisphere or compensatory recruitment of right hemisphere homologues of left hemisphere speech areas. Since this overactivity in the right hemisphere remains during both covert speech and covert humming conditions, it may not reflect reduced left-lateralised activity for language functions but instead an inhibitory response present in adults who stutter. On the other hand, it has been reported that certain therapeutic interventions for stuttering have demonstrated the potential to enhance neural activity within the left hemisphere of the brain (Toyomura et al., 2015) or shift the balance of activity from the right hemisphere to the left during speech production (Kell et al., 2009; Neumann et al., 2005). However, neither of these studies statistically compared the activity between two hemispheres in PWS and controls, which may explain why our results differ from these previous studies that we did not find a difference in laterality. As discussed above, overactivation in one hemisphere during one task could occur for various reasons, such as inhibition, compensation or error responses. Furthermore, the rightwards overactivity may be due to statistical thresholding, giving the impression that there is no activity in one hemisphere because it is only visible sub-threshold.

To our knowledge, there is only one study that found support for the idea that PWS are differently lateralized for language, which also investigated language laterality directly. Using near-infrared spectroscopy, Sato et al. (2011) reported laterality during passive listening of syllable pairs, which included either a phonemic contrast (e.g., /itta/ vs. /itte/) or prosodic contrast (e.g., /itta/ vs. /itta?/). Specifically, adults, school-aged children and pre-school children who stutter did not show the expected leftward lateralisation for processing the phonemic contrast, whereas controls showed a pattern of leftward and rightward lateralisation for phonemic and prosodic contrasts, respectively (Sato et al., 2011). Although the results of a direct comparison of language laterality between PWS and controls differ from our findings, it must be considered that this study involved passive listening to words rather than production.

Studies on structural hemispheric differences in PWS can offer valuable insights for interpreting functional activity. For instance, some earlier structural imaging studies in PWS reported lower leftwards (or more rightwards) asymmetry in the planum temporale, which was correlated with stuttering severity (Foundas et al., 2004). However, analysis of planum temporale asymmetry with a large population overlapping with the participants reported here found no differences between PWS and TFS and also no relationship with handedness (Gough et al., 2018).

### 4.2. LIs are More Strongly Left Lateralised for the Covert Tasks

In our findings, covert language tasks were significantly more lateralised compared with overt tasks. This finding is consistent with another fMRI study that reported more robust language lateralisation for covert language tasks than overt ones (Partovi et al., 2012). The reason may be that the cortical motor areas that send hundreds of commands to dozens of muscles bilaterally during overt speech production are not involved in covert speech. When the motor cortex is heavily involved in overt articulation, perhaps this bilateral pattern of task-related activity reduces laterality measured by methods that include these areas. As seen in our overt tasks (see Fig. 1), pre- and post-central gyri are reliably activated bilaterally during overt speaking tasks compared with a passive baseline task (meta-analysis from PET and fMRI studies: Turkeltaub et al., 2002). It is worth noting, however, that the analysis of the activity in BA44 and BA45 regions of interest, that do not include data from the pre- and post-central gyri, also show stronger lateralisation for the covert than the overt tasks (see Supplementary figures).

Another possible reason for stronger laterality during covert tasks in our findings needs to be highlighted. Both tasks (covert sentence reading and auditory naming) involved continuous data acquisition during imaging. In contrast, the overt speech production tasks were carried out using sparse sampling to allow participants to hear themselves during the task and to avoid potential head movement artefacts due to speech production during acquisition of the imaging data. Sparse sampling is feasible because the haemodynamic response is slow, only peaking some 4-6 seconds after the event, hence speech production can occur during a relatively long window of silence (6-10 s across different studies) before the measurement that takes 2-3 s occurs (Hall et al., 1999). Continuous imaging is debatably more sensitive since more data is sampled relative to sparse sampling, which only acquires a single measurement timed to coincide with the peak of the haemodynamic response evoked by the task. It is unclear why laterality should be different under these different measurement conditions since they would affect both hemispheres equally. Nevertheless, with our current datasets, we cannot disentangle possible causes of our finding that covert tasks were more strongly lateralized than overt ones since this factor is confounded with the measurement difference.

### 4.3. Limitations

Laterality indices allow us to compare the activity of thousands of voxels between two hemispheres in a single metric; however, useful information may be lost with this approach. LI scores deriving from fMRI results are limited in temporal resolution, which is not informative for possible differences in lateralisation milliseconds before the speech starts. For example, Neef and colleagues (2015) used transcranial magnetic stimulation during a verb generation task to record motor-evoked potentials from the tongue. They reported atypical cortical excitability patterns between left and right tongue motor cortical areas in adults who stutter milliseconds before speech onset. The current fMRI study would not be able to detect these dynamic differences in cortical excitability because of the relatively slow signal of the haemodynamic response (Glover, 2011).

In comparing activity between hemispheres, choices need to be made regarding the regions of interest. The advantage of large regions of interest, such as whole hemispheres or lobes, is that spatial variation in language processing among participants will be captured. The disadvantage of large regions is that many of the voxels included do not show task-evoked activity. ROI selections were made relying on masks of the frontal and temporal lobes, as these encompass the areas where language tasks usually elicit activity. In addition, we quantified the LI values within more focal regions of interest, specifically the pars opercularis (BA44) and pars triangularis (BA45) situated within the inferior frontal gyrus, thereby affording a finer-grained examination.

## 5. Conclusions

Previous fMRI findings consistently reported an overactive right hemisphere in stuttering during speech tasks but did not statistically compare the functional activity between hemispheres. Therefore, they do not provide direct evidence for altered hemispheric specialisation in people who stutter during language production. Here, our results close that gap by statistically comparing the functions of two hemispheres to test the altered hemispheric specialisation theory with a threshold-independent laterality analysis. Our findings indicated no difference in the hemispheric specialisation in frontal and temporal regions of PWS compared with typically fluent speakers while performing four different speech and language tasks. We also reported that covert tasks were substantially more lateralised than overt tasks for both groups. These data were obtained using continuous imaging, whereas the overt tasks were carried out using sparse sampling. Therefore, task choice and data acquisition may be important factors to consider when measuring laterality.

## Supporting information

Supplementary Table Scan Details

## Data Availability

The datasets analysed during the current study are available in the OSF (doi: 10.17605/OSF.IO/TQ8J5). This includes laterality indices, demographics, and stuttering severity scores for each participant. The brain imaging data of the group averages in the standard space are available at NeuroVault: https://neurovault.org/collections/16872/.

## Abbreviations

PWS: people who stutter
TFS: typically fluent speakers
LI: laterality index
ROI: region of interest.

## Acknowledgements

We sincerely thank the Dominic Barker Trust for their support of our research. We also express our gratitude to Dr. Zoe V. J. Woodhead for her assistance with the laterality analysis. Our sincerest thanks go to all the participants who took part in this study. For the purpose of open access, the author has applied a CC BY public copyright licence to any Author Accepted Manuscript version arising from this submission.

## Funding

This research was supported by the Dominic Barker Trust (Registered Charity No. 1063491) and NIHR Oxford Health Biomedical Research Centre (NIHR203316). The views expressed are those of the author(s) and not necessarily those of the NIHR or the Department of Health and Social Care. The Wellcome Centre for Integrative Neuroimaging is supported by core funding from the Wellcome Trust (203139/Z/16/Z and 203139/A/16/Z). The original funding for the data collection came from: grants to KW from the Medical Research Council (G0400298) and a British Academy/Leverhulme Trust small project grant (SG130103); a Clinical Research Training Fellowship to JC from the Medical Research Council (MR/K023772/1); and a Stammer Trust grant and a Reading University Functional Imaging Facility New Directions grant to DW.

## Notes

https://osf.io/tq8j5/?view_only=1bab145f1632444eaea5590bb18ed06b

### Competing Interest Statement

The authors have declared no competing interest.

