## Supplementary Figures BA44 & BA45 for "No Evidence of Altered Language Laterality in People Who Stutter across Different Brain Imaging Studies of Speech and Language"

### LIs for task-evoked activity in Pars Opercularis (BA44)

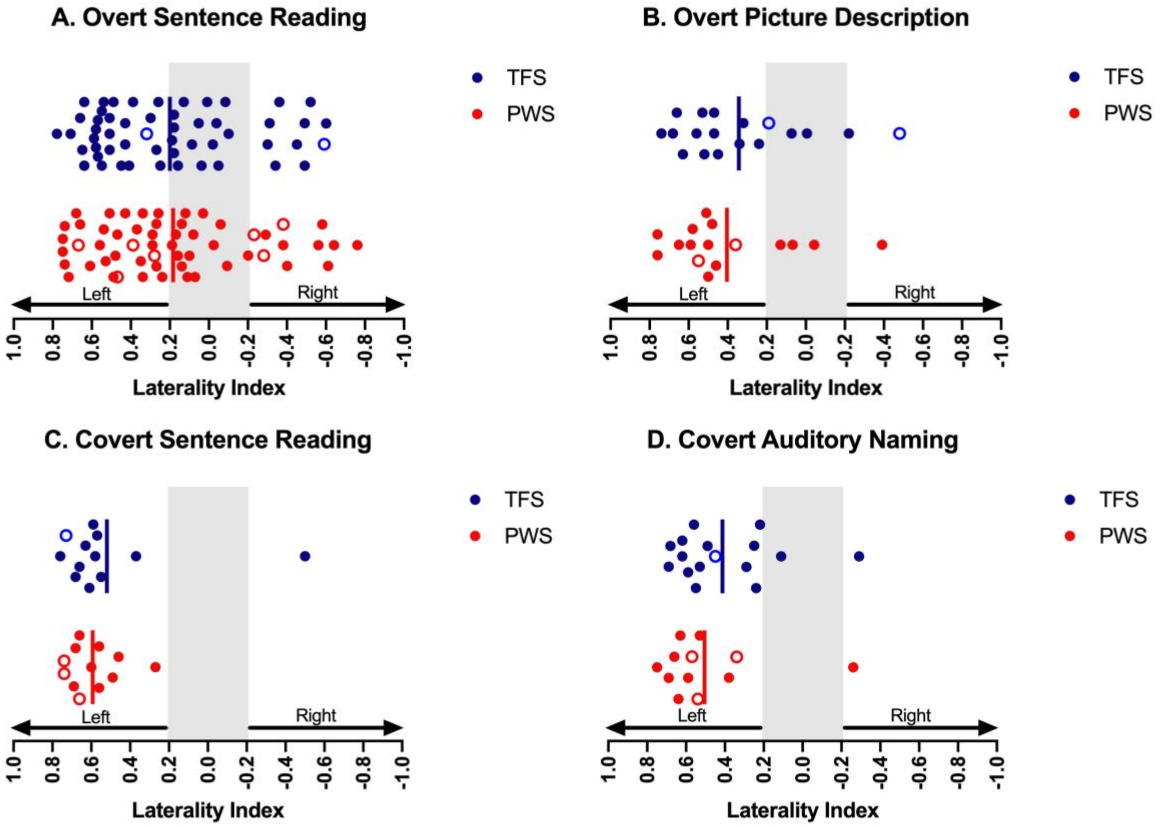


**Figure 1: Laterality indices in people who stutter (PWS) and typically fluent speakers (TFS) based on activity in the frontal lobes.** Solid vertical lines represent group means. The grey area represents LI values between -0.2 and 0.2, which are considered not lateralised. PWS: People who stutter are the red circles; TFS: Typically fluent speakers are the blue circles. The circles indicate left-handedness in both groups.

### LIs for task-evoked activity in Pars Triangularis (BA45)

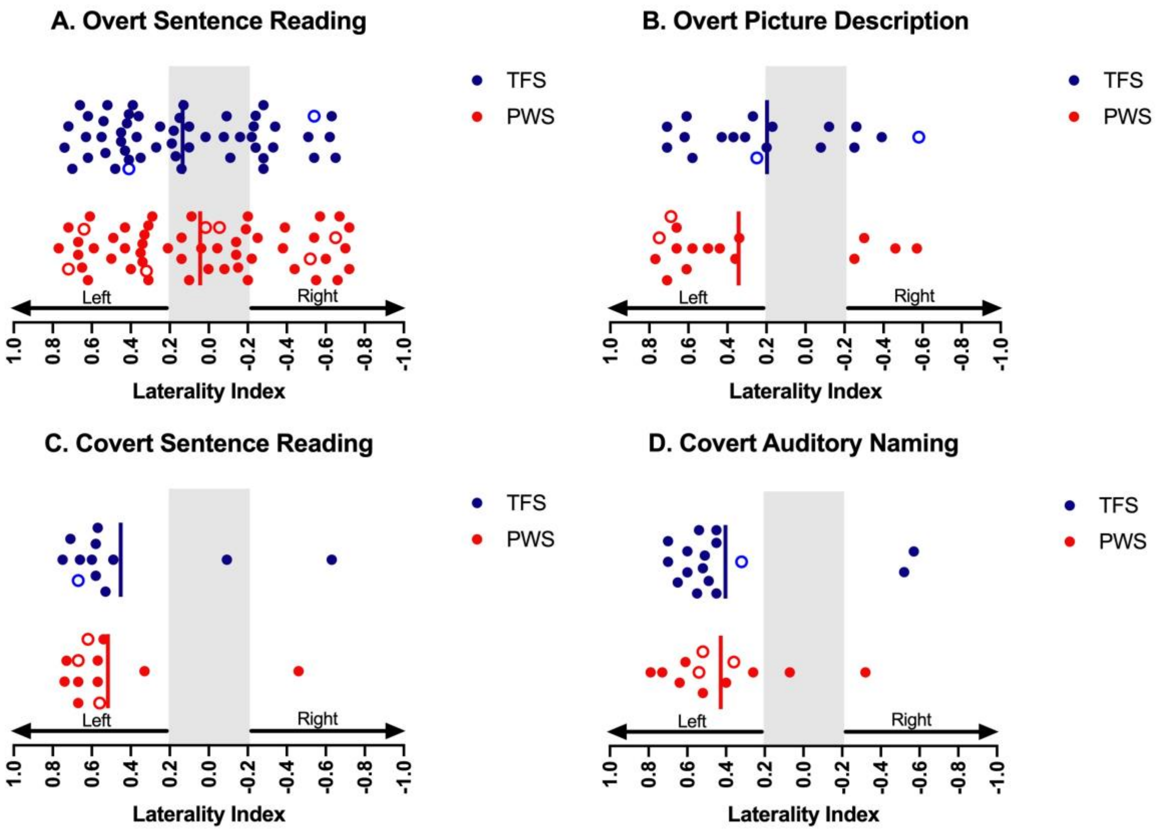


**Figure 2: Laterality indices in people who stutter and typically fluent speakers based on activity in the temporal lobes (see Figure 1 for details).**
