## Supplementary Table Scan Details for "No Evidence of Altered Language Laterality in People Who Stutter across Different Brain Imaging Studies of Speech and Language"

| Task | Baseline | PWS | CON | Scanner | TE (ms) | TR/TA (s) | Slices (N) | Thick-ness (mm) | In-plane (mm) |
|---|---|---|---|---|---|---|---|---|---|
| Overt sentence reading (NAF/DAF)[1] | Row Xs | 10 | 10 | 3T Varian-Siemens | 30 | 10 / 3 | 32 | 4 | 4 x 4 |
| Overt sentence reading (NAF/DAF)[2] | Row Xs | 8 | 11 | 3T Siemens Trio (1) | 30 | 10 / 3 | 50 | 3 | 3 x 3 |
| Overt sentence reading (NAF)[3] | Farsi script | 23 | 15 | 3T Siemens Trio (1) | 30 | 9 / 2 | 38 | 3.5 | 3 x 3 |
| Overt sentence reading / Picture description[4] | Fixation | 16 | 18 | 3T Siemens Trio (2) | 30 | 9 / 2 | 32 | 4 | 3 x 3 |
| Covert Sentence Reading / Covert Reading & listening / Passive Listening[2,5] | Row Xs | 12 | 12 | 3T Siemens Trio (1) | 30 | 2.4 | 40 | 3 | 3 x 3 |
| Covert Auditory Naming[2,6] | Reversed Speech / Fixation | 12 | 16 | 1.5T Siemens Sonata | 50 | 3 | 35 | 4 | 3 x 3 |

[1] Watkins et al., 2008

[2] Unpublished data

[3] Chesters et al., 2021 (preprint)

[4] Connally et al., 2018

[5] Erb et al., 2010 (abstract)

[6] Paradigm described in Badcock et al., 2012
